# SCsnvcna: Integrating SNVs and CNAs on a phylogenetic tree from single-cell DNA sequencing data

**DOI:** 10.1101/2022.08.26.505465

**Authors:** Liting Zhang, Hank W. Bass, Jerome Irianto, Xian Mallory

## Abstract

Single-cell DNA sequencing enables the construction of evolutionary trees that can reveal how tumors gain mutations and grow. Different whole genome amplification (WGA) procedures render genomic materials of different characteristics, often suitable for the detection of either single nucleotide variation (SNV) or for copy number aberration (CNA), but not for both, hindering the placement of both SNVs and CNAs on the same phylogenetic tree for the study of interplay of SNVs and CNAs. SCARLET places SNVs on a CNA tree, a tree derived based on the copy number profiles, while considering SNV loss due to copy number losses. However, SCARLET requires that the SNVs and CNAs are detected from the same sets of cells, which is technically challenging due to the sequencing errors or the low sequencing coverage associated with a particular WGA procedure. Here we presented a novel computational tool, SCsnvcna, that aims at placing SNVs on a CNA tree whereas the sets of cells rendering the SNVs and CNAs are independent, thus is more practical in terms of the technical challenge from single cell WGA process. SCsnvcna is a Bayesian probabilistic model that utilizes both the genotype constraints on the tree and the cellular prevalence (CP) to search the solution that has the highest joint probability. Both simulated and real datasets show that SCsnvcna is highly accurate in predicting the placement of SNVs and SNV cells. In addition, SCsnvcna has a precise prediction of SNV losses due to copy number loss.

## Introduction

Cancers result from a combination of abnormal cell growth along with metastasis, the ability to spread to other tissues. The malignant progression of cancer is associated with acquired mutations that either activate oncogenes or inhibit tumor suppressor genes [1,10,12]. Copy number aberrations (CNA) [4] and single nucleotide variants (SNV) [1] are two major types of acquired mutations that lead to cancer, yet these two differ in mechanism and scale (single base pair vs. regions of various sizes), and they are detected using different technologies.

During the growth, cancer cells continue to acquire and pass on different mutations, leading to the Intra Tumor Heterogeneity (ITH) problem [7, 14, 21]. ITH explains how different cells from the same cancer can have different genotypes or mutational spectra, resulting in heterogeneous cellular phenotypes with differential response to therapies, including drug resistant cancer cells [7, 20]. ITH confounds both diagnosis and treatment strategies. To better understand ITH, it is essential to be able to categorize the group of mutations in different lineages or subclones of the cancer, and reconstruct the mutational lineages by combining information from SNV and CNA approaches. For example, the “two-hit” hypothesis assumes a loss of an allele in a suppressor gene followed by a gain of mutation on the remaining allele [12]. However, the “bulk sequencing”, which combines all sampled cells together for sequencing, does not differentiate subclones from each other directly [31].

Single-cell DNA sequencing (scDNA-seq) technologies makes it possible to decipher ITH by sequencing one cell at a time [22]. However, due to the limited quantity of about 6pg genomic DNA in a normal human somatic cell, scDNAseq typically requires whole genome amplification (WGA) before the library construction [3], and different WGA procedures lead to data with different error profiles. The two most popular scDNA-seq WGA techniques are multiple displacement amplification (MDA) [11, 29] and degenerate oligonucleotide-primed PCR (DOP-PCR) [2, 5, 22]. The MDA method gives a higher genome recovery, suitable for SNV detection. However, the uneven genome amplification from the MDA makes it not suitable for CNA detection [19]. DOP-PCR, by contrast, generates relatively uniform coverage of reads, suitable for large-region CNA detection [19]. However, DOP-PCR has a lower proportion of genome coverage, limiting its utility for detection of SNVs from a single cell [29].

So far there has been no WGA technique reliable for the simultaneous detection of CNA and SNV from single cells. The two WGA methods require that the DNA from any single cell be used for CNA or SNV, but not both. Consequently, single-cell SNV and CNA assays can be done in parallel from a common sample, but produce separate datasets from two different sets of cells. This leads to two phylogenetic trees, one from CNAs and the other from SNVs. Should we be able to integrate or combine information from these two phylogenetic trees into one composite tree, we could recover a more complete biological evolutionary tree. This would create new opportunities to investigate the relationship among SNVs and CNAs in terms of their interplay and order with each other within a single tumor sample, which can further help us characterize ITH.

Unfortunately, so far there has been no published tool that can integrate the SNVs and CNAs on a phylogenetic tree given the scDNA-seq data prepared by different WGAs prior to library preparation. PACTION [24] integrates both SNV and CNA signals in identification of tumor clones, but it was not designed for scDNAseq data. Scarlet [25] constructs tumor phylogenies from both CNAs and SNVs and supports the loss of SNVs due to the copy number losses. However, it requires that CNAs and SNVs to be present on the same set of cells. Due to the read depth nonuniformity on the sequences of SNV cells, the detection of CNAs has to be on a very large scale, leading to a low-resolution CNA tree to start with. BiTSC^2^ [6] and COMPASS [27] are the other two methods that jointly infer a phylogenetic tree from both SNVs and CNAs. However, like Scarlet, they also require that CNAs and SNVs are from the same set of cells. Moreover, BiTSC^2^ does not explicitly output the placement of mutations on the tree whereas COMPASS is designed mainly for Mission Bio Tapestri platform data which has been used for targeted PCR and have limited coverage on the genome. Phertilizer [30] infers a phylogenetic tree with both SNVs and CNAs, but the SNVs are inferred from the cells sequenced mainly for CNA detection, leading to a high missing rate in SNVs. Although Phertilizer tries to increase the detection rate of SNVs by inferring the clones of the cells, the SNV detection rate depends heavily on the size of the clones, the sequencing coverage of the cells and the number of cells being sequenced. Dorri et al. [8] infers the phylogenetic tree based on CNAs and can infer the SNVs based on the inferred tree. However, like Phertilizer, method proposed by Dorri et al. infers CNA and SNV from the same set of the cells, leading to their SNV calling suffering from high missing rate while constrained to clone size, sequence coverage and the number of sequenced cells. Leung [15] et al. paired up clones of SNV cells and CNA cells and annotated CNAs on a SNV tree on two metastatic colorectal cancer samples, but such integration was performed mostly manually, and it was on a low resolution instead of on the cell granularity, requiring the clustering of the cells prior to the integration.

Since scDNA-seq is essential in deciphering ITH, we developed SCsnvcna, a publicly available computational phylogenetics tool that combines SNVs and CNAs from independent single-cells data sets of the same sample into one phylogenetic tree using a Bayesian probabilistic model. SCsnvcna has the following features that are advantageous when applied to scDNA-seq.

– SCsnvcna is error-aware. MDA typically produces data that has a high false positive (FP) and false negative (FN) SNVs [32] in addition to the missing data due to the lack of coverage. SCsnvcna models all three types of errors to improve the accuracy of error predictions.
– SCsnvcna is bias-aware. SCsnvcna models the cell sampling bias when the percentage of the cells carrying SNVs does not agree with the percentage of the cells carrying CNAs occurring on the same tree branch. Furthermore, it can also predict the bias.
– SCsnvcna can overcome the loss of SNVs due to copy number losses, and thus implements the loss-supported evolutionary model proposed by SCARLET [25].

## Methods

### Model Description

SCsnvcna requires two inputs, a ternary matrix *D*_*n∗m*_ and a phylogenetic tree. *D*_*n∗m*_ is the observed binary genotypes for SNVs inferred from scDNA-seq data, where *n* is the number of the single cells suitable for SNV detection (called **SNV cells** in the following text) and *m* is the total number of SNV sites. Each entry *D*_*i,j*_ *∈* [0, 1, *X*], where *i* = 0, 1, …, *m* and *j* = 0, 1, …, *n*. Here the entry values 0, 1 and *X* denote the absence, the presence and the missing entry for SNV cell *i* at SNV site *j*. The second input is a phylogenetic tree *T* inferred from CNA signals. We call such a tree inferred purely from CNA signals a **CNA tree**. *T* is a rooted directed binary tree with *K* edges and *K* + 1 nodes. The root of the tree represents a normal cell that has no somatic CNAs or SNVs. The leaves of the tree represent the observed single cells that have the copy number signal, for example, the cells sequenced from DOP-PCR library preparation. We call these cells suitable for CNA detection **CNA cells**. The internal nodes represent ancestral cells that are not observed in the data. The CNA tree *T* can be obtained by running existing tools such as PAUP [28] or SCICoNE [13].

We assume that *D*_*n∗m*_ and *T* are observed from the same patient sample, and the SNV cells and CNA cells are randomly sampled from the same cell pool. The major goal of SCsnvcna is to place SNVs on the edges of the CNA tree and predict the error rates.

To achieve this goal, SCsnvcna uses a probabilistic model and Monte-Carlo Markov Chain (MCMC) to sample the placement of the SNVs on the edges of the CNA tree and the hyperparameters *G, θ* and *σ*, in which *G* is the underlying genotype matrix, *θ* is a set of SNV cell error rates including FP rate *α*, FN rate *β* and missing rate *γ*, and *σ* models the difference of the cellular prevalence between SNV cells and CNA cells. **Fig. 1** gives an illustration of the model. In particular, given the input matrix *D* and CNA tree, the MCMC searches for the true underlying *G* matrix, the placement of SNVs, the placement of SNV cells and the placement of the hyperparameters *θ* and *σ* in an iterative fashion. In the following text, we describe in more detail the incorporation of single cell errors (*θ*), the CP variance between CNA and SNV cells (*σ*) and the consideration of mutation loss due to copy number loss.

**Fig. 1:**
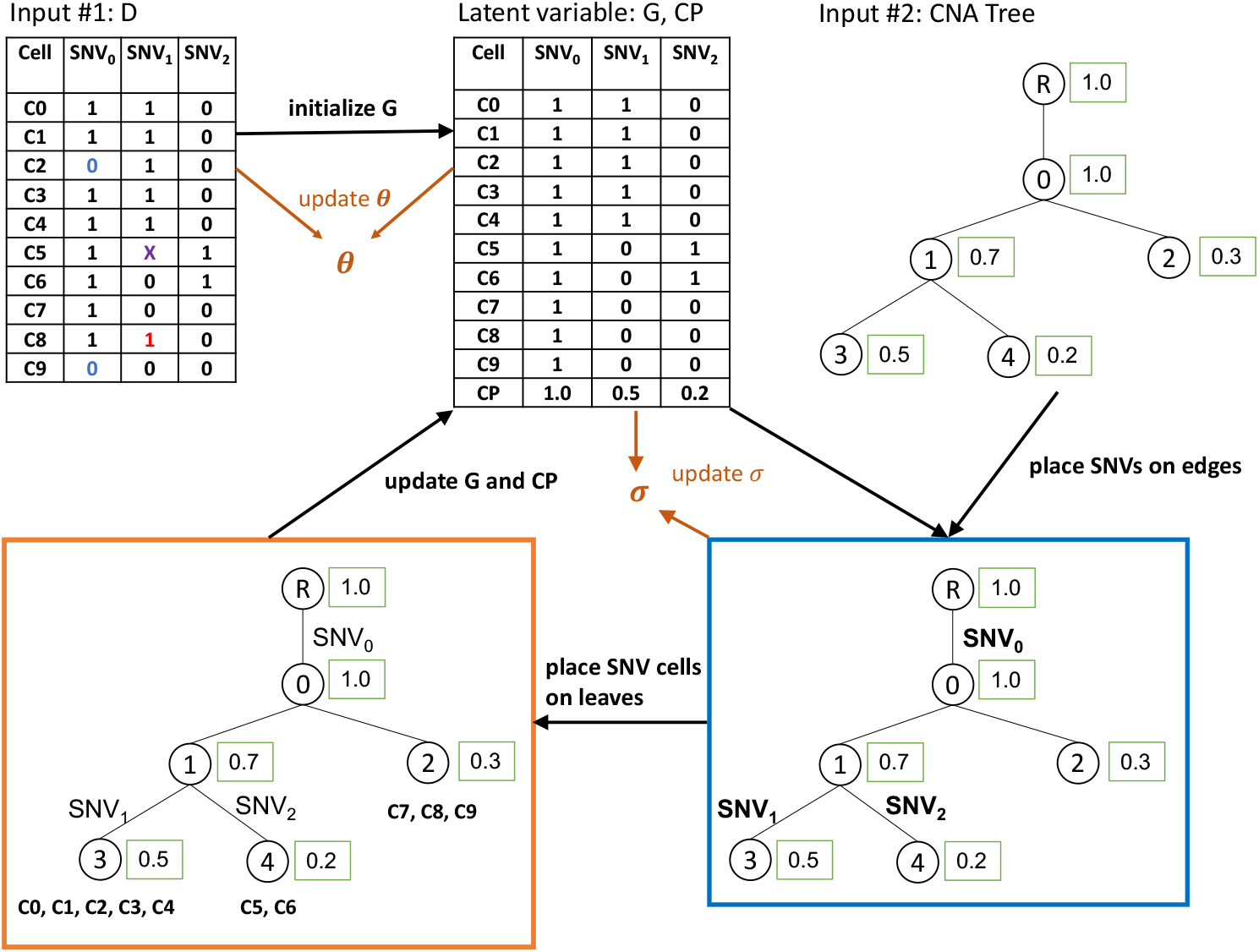
Illustration of SCsnvcna method. Inputs are observed data *D* (a cell by SNV matrix) and a CNA tree. Observed data may have false positive SNVs (red “1”s), false negative SNVs (blue “0”s) and missing entries (“X”). The underlying genotype matrix *G* along with the cellular prevalence (CP) are initialized based on *D*. SCsnvcna places SNVs on the edges of the CNA tree (blue box), SNV cells on the leaves of the CNA tree (orange box), followed by the update of the latent variables which are *G*, the error rates *θ* and the CP variance variable *σ*. The output of SCsnvcna is the placement of SNVs and cells on the CNA tree, the predicted values of *G, θ* and *σ*.

### Model of single-cell errors

The observed genotype matrix of SNVs *D* is different from the true underlying genotype matrix *G*, a *n ∗ m* binary genotype matrix, due to the sequencing errors. We model the errors in *D* as *θ*, where *θ* = { *α, β, γ}*, in which *α, β* and *γ* represent FP, FN and missing error rates. The probability of *D* given *G* and *θ* is described in Eq. 1.

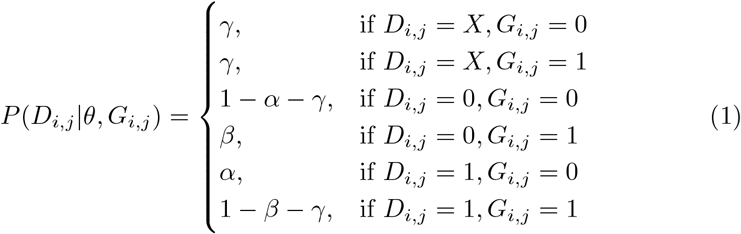

### Using the cellular prevalence (CP) to place SNVs on a CNA tree

The CP of a SNV *j*, 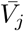, can be calculated by Eq. 2 given its true underlying genotype *G*_*i,j*_ for all cells.

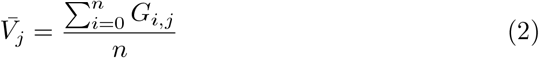

Given the CP for a SNV *j*, SCsnvcna searches the most fitting branch on tree *T* to place SNV *j*. Specifically, suppose the percentage of CNA cells in *T* under an edge *E*_*k*_ is 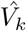. We place SNV *j* on the edge *E*_*k*_ whose 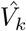 is closest to 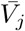. This is based on the assumption that SNV and CNA cells are from the same patient sample and thus the same underlying phylogenetic tree. To consider potential sampling bias, we model the distance between the two CPs, 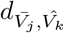 by a Gaussian distribution which has zero mean and *σ* as the standard deviation. *σ* represents the sampling bias which is a hyperparameter to be inferred by our MCMC sampling algorithm or given by the user.

### Using the CNAs on SNV cells to limit the placement of SNV cells

Although the SNV cells are mainly for the detection of SNVs, it is still possible to detect some CNAs at a lower resolution. Should these CNAs overlap with those CNAs detected from CNA cells, SCsnvcna will constrain the SNV cells that contain these CNAs to the leaves whose CNA cells contain the same CNAs. Since the path from the root to the node where the SNV cell is placed spells out the SNVs that the cell carries, the placement of SNV cells constrains the placement SNVs. Thus the CNA signal on SNV cells, although limited, constrains not only the placement of SNV cells, but also the placement of SNVs on the edges of the CNA tree.

Deciding which leaves on the CNA tree that the SNV cells are constrained to is a preprocessing step of SCsnvcna. We implemented two methods to provide a list of leaf nodes and users can choose either way should they provide the copy number profiles of both CNA cells and SNV cells. The first method is applicable when there are very limited number of CNAs detectable on the SNV cells. Specifically, for a SNV cell *i*, if one of its CNAs overlaps with a CNA *u* detected from CNA cells by at least *p*%, we limit the placement of the SNV cell, *i*, on only the leaves of the subtree whose root is the node below the edge where *u* occurs. If more than one CNA on the SNV cell *i* overlap with those detected from CNA cells, the placement of cell *i* will be the intersection of the nodes constrained by each of the overlapping CNAs. Unlike the first method, the second method compares pairwisely the entire copy number profiles between all CNA cells and all SNV cells. If a SNV cell and a CNA cell have ≥ *q*% of overlapping CNAs, the SNV cells’ placement is restricted to the node where the CNA cell is placed. In our real data analysis, since the SNV cells are informative of the CNAs, we applied the second method in analyzing the real data sample coming from a scDNA-seq study of metastatic colorectal cancer (CRC2) [14].

### Loss of SNVs due to copy number loss

It is possible that a SNV is lost due to the copy number loss. Thus all the SNV cells that obtained the SNV followed by a copy number loss may not show signal of this SNV on the *D* matrix any more. SCsnvcna is capable of detecting such SNV loss due to copy number loss. Should SCsnvcna determine a certain SNV is lost due to copy number loss, there is no penalty in the CP distance between the CNA cells and SNV cells, i.e., the nodes under the edge of the copy number loss are not counted in calculating the CPs of CNA cells or SNV cells.

In the case of multiple equally good solutions while considering the loss of SNVs due to copy number loss, we randomly select one of the best trees.

### Probabilistic graphic model of SCsnvcna Fig. 2

shows the probabilistic graphic model of SCsnvcna.

**Fig. 2:**
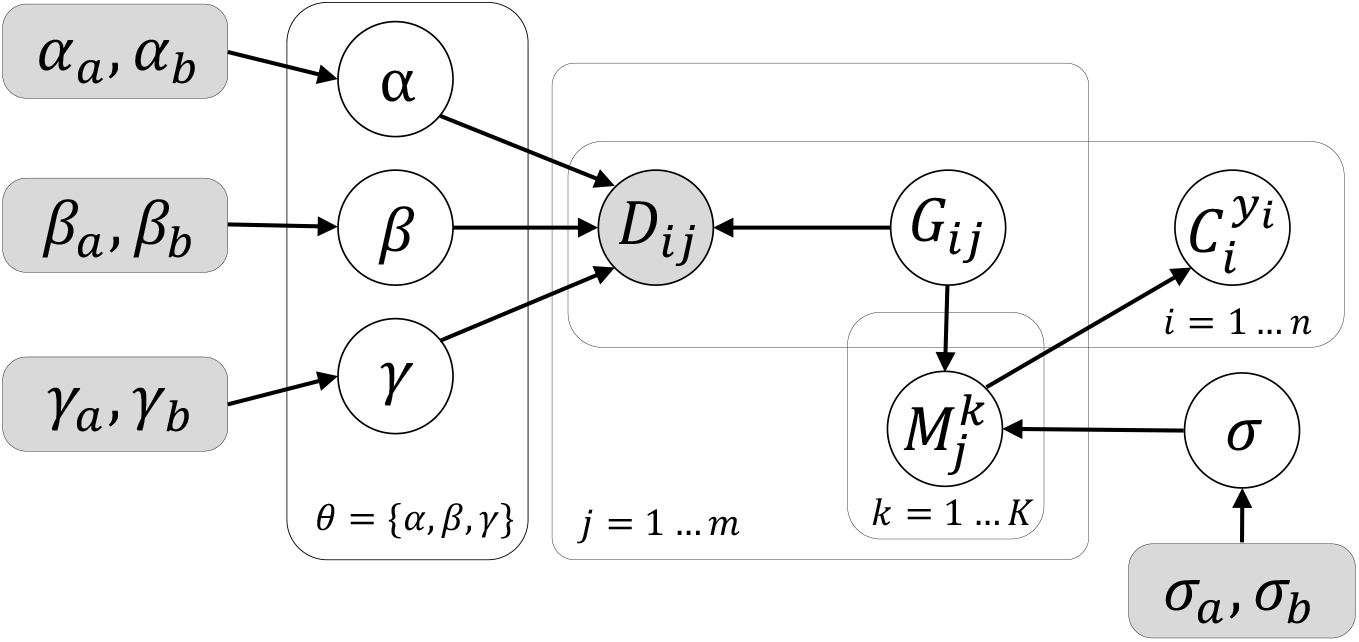
Probabilistic graphic model of SCsnvcna. Shaded nodes are given parameters or input data. Unshaded nodes are the latent variables to be inferred.

The distributional assumptions for all variables are as follows.

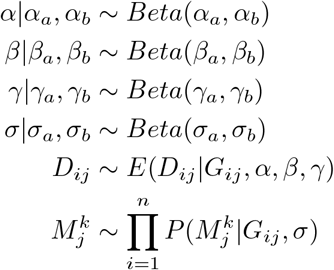

where *E* denotes the error model in the data defined by Eq. 1. *α*_*a*_, *α*_*b*_, *β*_*a*_, *β*_*b*_, *γ*_*a*_, *γ*_*b*_, *σ*_*a*_ and *σ*_*b*_ are hyperparameters controlling *α, β, γ*, and *σ*. 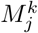 denotes the event of placing SNV *j* on edge *E*_*k*_. 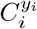 denotes the event of placing cell *i* on leaf *y*_*i*_. 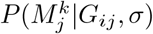 denotes the probability that SNV *j* occurs on edge *E*_*k*_ given all the cell placement and correct genotypes. 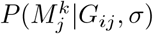 can be rewritten as 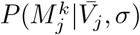 as we use the CP to find the best fitting placement of a SNV whereas 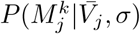is defined in Eq. 3.

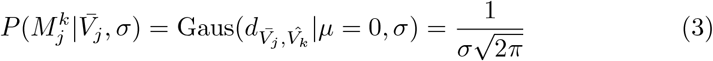

To consider the loss of SNVs due to copy number loss, when a SNV overlaps with a CNA, we define 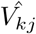 as the fraction of the CNA cells that share edge *E*_*k*_ on the path to the root and have no copy number losses overlapping with mutation *j*. 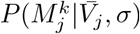 then becomes

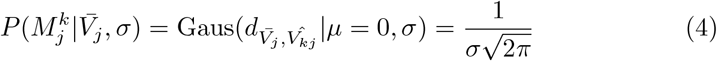

We use Eq. 4 to calculate 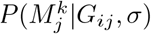 to facilitate the detection of mutation losses due to the copy number losses.

The joint distribution of all parameters is

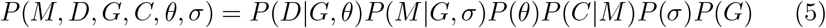

### MCMC sampling

SCsnvcna’s MCMC sampling is composed of three parts: 1) the placement of SNVs which is coupled with the sampling of the placement of SNV cells, 2) *θ* and 3) *σ*. For 1), we first make a proposal of the change of a SNV placement on any edge other than the current one that the SNV is placed. To decide whether we will accept such a replacement, we consider the distances of the CPs between the SNV cells and CNA cells as well as the error model of SNVs cells given the previously updated *G* and *θ*. The calculation of these probabilities requires the updated placement of cells corresponding to the new placement of SNVs. We therefore use the maximum likelihood to place the cells so that the updated placement of cells can best explain data *D* given the updated SNV placement, and previously updated *G* and *θ*. To be more specific, the probability of placing cell *i* on leaf *y*_*i*_ given that SNV *j* comes from branch 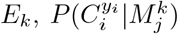, can be redefined as 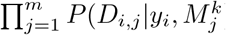, whereas 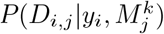 is defined in Eq. 6.

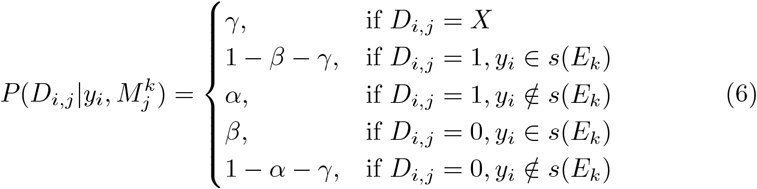

where *s*(*E*_*k*_) is the set of leaves who share the branch *E*_*k*_ on the path to the root.

To model the loss of SNVs due to loss of a copy number, the distance between CPs is only calculated on the nodes that are not subject to the mutation loss due to copy number loss. Thus Eq. 6 is modified to be Eq. 7 when considering potential mutation losses.

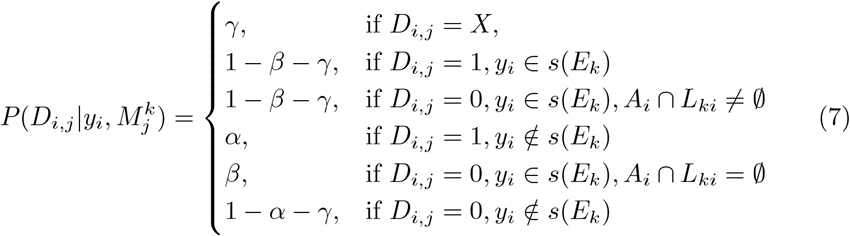

in which *A*_*i*_ are the set of the copy number losses that mutation *i* overlaps with, and *L*_*ki*_ are the set of the copy number losses imputed on the edges along the path from the edge *E*_*k*_ to leaf *y*_*i*_.

The updated placement of SNVs and cells allows the update of *G* and 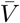 so that each cell’s SNVs are consistent with the new placements of SNVs and cells. The Metropolis-Hastings acceptance ratio of the proposed SNV placement is calculated by Eq. 8.

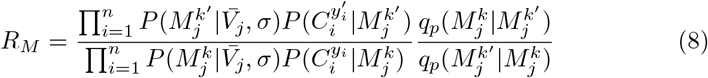

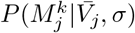 and 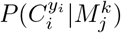 can be calculated by Eq. 3 and Eq. 7, respectively.

The second sampling is for a new set of *θ*. In this step, we propose new *θ*^*′*^ = { *α*^*′*^, *β*^*′*^, *γ*^*′*^*}* given the current error rate by a Gaussian distribution. For each parameter in *θ*, we fix the current value as mean and 1 as the standard deviation in the Gaussian distribution. The Metropolis-Hastings acceptance ratio for *θ*^*′*^ is given by Eq. 9.

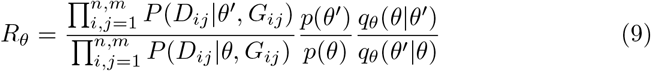

Notice that we sample *α, β* and *γ* one at a time but use *θ* to summarize all three of them in Eq. 9.

To effectively sample a solution and reduce the running time, we constrain our *α* to ≤ 0.1 and *β* to ≤ 0.6 and thus reject all the samplings which are not in the above-mentioned range. These constraints are applied due to their consistency with the biological prior knowledge of these two error rates [32].

Without the loss of generality, the three parameters are sampled in the order of *α, β, γ*.

The third sampling is for a new *σ*. The proposed *σ*^*′*^ is sampled from a Gaussian distribution whose mean is the current *σ* and standard deviation is 1. The Metropolis-Hastings acceptance ratio for the proposed *σ*^*′*^ given the current *σ* is described in Eq 10.

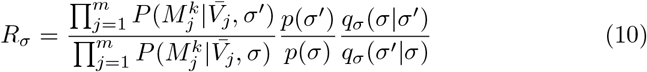

We summarized the MCMC sampling algorithm in **Supplemental Algorithm 1**.

### Initialization

#### Initializing *G*

For the sake of faster convergence of MCMC, we initialize *G* given the observed data *D*_*i,j*_ by Eq. 11.

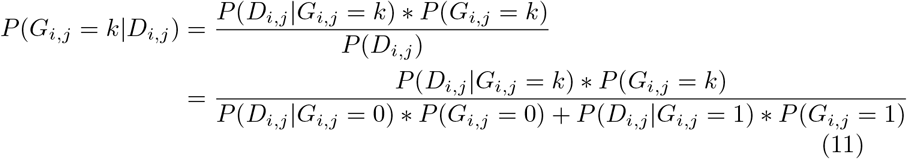

where *k* ∈ {0, 1}. *P* (*D*_*i,j*_|*G*_*i,j*_) is defined in Eq. 1, and *P* (*G*_*i,j*_) is the prior probability for *G*. The prior probability *P* (*G*_*i,j*_ = 1) is defined based on the estimation of the FP and FN rates as well as *D*. In specific, the probability *P* (*G*_*i,j*_ = 1) is defined such that the average value of *G* reflects the initialized value of *α* and *β* as well as *D*. In more details, suppose there are *t* 1’s and *u* 0’s observed in *D*, and *α*_0_ and *β*_0_ are the initial average FP and FN rates. We define *t*^*′*^ = *α*_0_ ∗ *t* and *u*^*′*^ = *β*_0_ ∗ *u* as the number of FP and FN entries in *D*, respectively. For each entry *i, j*, the prior probability *P* (*G*_*i,j*_ = 1) can be calculated as 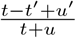.

#### Initializing hyperparameters *α*_*a*_, *α*_*b*_, *β*_*a*_, *β*_*b*_, *γ*_*a*_, *γ*_*b*_, *σ* and the placement of SNVs on the CNA tree

*α*_*a*_ and *α*_*b*_ are the hyperparameters for the Beta distributions of false positive rate in the SNV cells. So are *β*_*a*_ and *β*_*b*_ for false negative rates in the SNV cells. We set up *α*_*a*_, *α*_*b*_, *β*_*a*_, *β*_*b*_ such that the resulting false positive and false negative rates’ mean equal to the initial input values which are 0.01 and 0.2, respectively, if not specified by the user. We further fix *α*_*a*_, *α*_*b*_, *β*_*a*_, *β*_*b*_ so that the standard deviation of the resulting Beta distributions is large yet the distribution curve is convex. Similarly, *γ*_*a*_ and *γ*_*b*_ are set up such that the average *γ* equals to the fraction of entries missing in the SNV matrix whereas the standard deviation of the resulting Beta distribution for *γ* is large.

We initialize the SNV placement according to Eq. 3. We do not take into consideration the mutation loss while initializing SNV placement. The initialization of SNV cell placement is done by maximizing 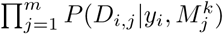 based on the initial placement of SNVs.

## Results

### Simulation

We developed a simulator described in **Supplementary Section 1** to comprehensively test SCsnvcna under different conditions. We varied nine variables in the simulation for testing SCsnvcna and compared it with the most state-of-the-art method, SCARLET [26], on these nine datasets. We did not compare SCsnvcna with Dorri, et al [8] and Phertilizer [30] mainly because like SCARLET, SCsnvcna infers the SNV placement on a CNA tree, assuming that the CNA tree is given, whereas Dorri, et al [8] and Phertilizer [30] focus on inferring a phylogenetic tree based on CNAs or both CNAs and SNVs. The nine variables are the number of subclones, false positive rates, false negative rates, missing rates, mutation loss rates, number of cells, number of mutations, a variable in the Beta splitting model for CNA tree generation, and lastly, the standard deviation of the CPs between SNV cells and CNA cells. The values used for each variable are listed in **Table S1**. For each dataset, we created the corresponding input files for SCARLET to make a fair comparison. We listed in more details the difference between SCsnvcna and SCARLET in **Supplementary Section 2.1**, and the whole process of simulating the input files for SCARLET and the comparison strategy between SCsnvcna and SCARLET in **Supplementary Section 2.2**.

For each dataset, we ran SCsnvcna with default parameter. In particular, MCMC search was run on 10000 iterations with 10 restarts and the solution with the highest posterior probability will be chosen as the final solution. For SCARLET, we ran it with default parameters.

#### Evaluation Strategy

We evaluate SCsnvcna’s performance by five metrics, 1) the error rate of the inferred underlying genotype matrix (we call it “genotype error”); 2) the error rate of the inferred ancestral relationship among SNV pairs (we call it “pairwise SNV error”); 3) the error rate of the inferred ancestral relationship between SNVs and CNAs (we call it “pairwise SNV and CNA error”); 4) sensitivity of mutation loss detection; and 5) specificity of mutation loss detection.

The first three metrics measure the accuracy of the placement of SNVs and SNV cells, whereas the last two metrics measure the accuracy of the detection of mutation losses.

On evaluating the ancestral relationship of a pair of SNV and CNA, we consider the following cases: 1) when a SNV and a CNA occur on the same edge; 2) when a SNV is ancestral to a CNA; 3) when a CNA is ancestral to a SNV; 4) a SNV and a CNA are incomparable. We apply the same four categories to the ancestral relationship of a pair of SNVs. If our inferred SNV placement leads to a different case from what was reported in the simulated ground truth tree, we count it as an error. We then report the average pairwise ancestral relationship error rate in the inferred SNV placement in terms of a pair of SNV and CNA, and a pair of SNVs.

Due to that SCARLET does not separate SNV cells and CNA cells as shown in **Supplementary Section 2.1**, for a fair comparison, we constrain our SNV cell placement only to the correct leaves in the simulation experiments.

We observe some cases when the lack of SNV cells loosens the constraints of SNV placements, thus a SNV can be placed equally well on multiple edges, an example seen in **Fig.S1**. Equally good placements of a SNV are the set of edges where the SNV can be placed without changing the resulting genotype matrix whereas one of the edge is the underlying true placement of the SNV. In our evaluation, all equally good placements of a SNV are counted as correct placements.

#### Simulation Results

We applied both SCsnvcna and SCARLET to the nine sets of simulated data and compared the five metrics of their results, as shown in **Fig. 3, Fig. 4, Fig.S2** and **Fig.S3**. In all the simulated datasets, SCsnvcna’s genotype error rate, pairwise SNV error rate and pairwise SNV and CNA error rate were much lower than that of SCARLET’s. Except the standard deviation of CPs and the Beta splitting variable whose results were shown in **Fig.S2**, we noticed a clear trend of increasing or decreasing error rate with the increasing or decreasing varying variable for all nine variables. Of the rest of the seven variables, we divided them into two groups. The first group focused on the parameters that may affect the accuracy of SCsnvcna and was composed of false positive rate, false negative rate, missing rate and mutation loss rate, the result of which were shown in **Fig. 3**. The second group focused on testing the scalability of SCsnvcna and was composed of the number of subclones in the tree, number of cells and number of mutations, the result of which was shown in **Fig. 4**.

**Fig. 3:**
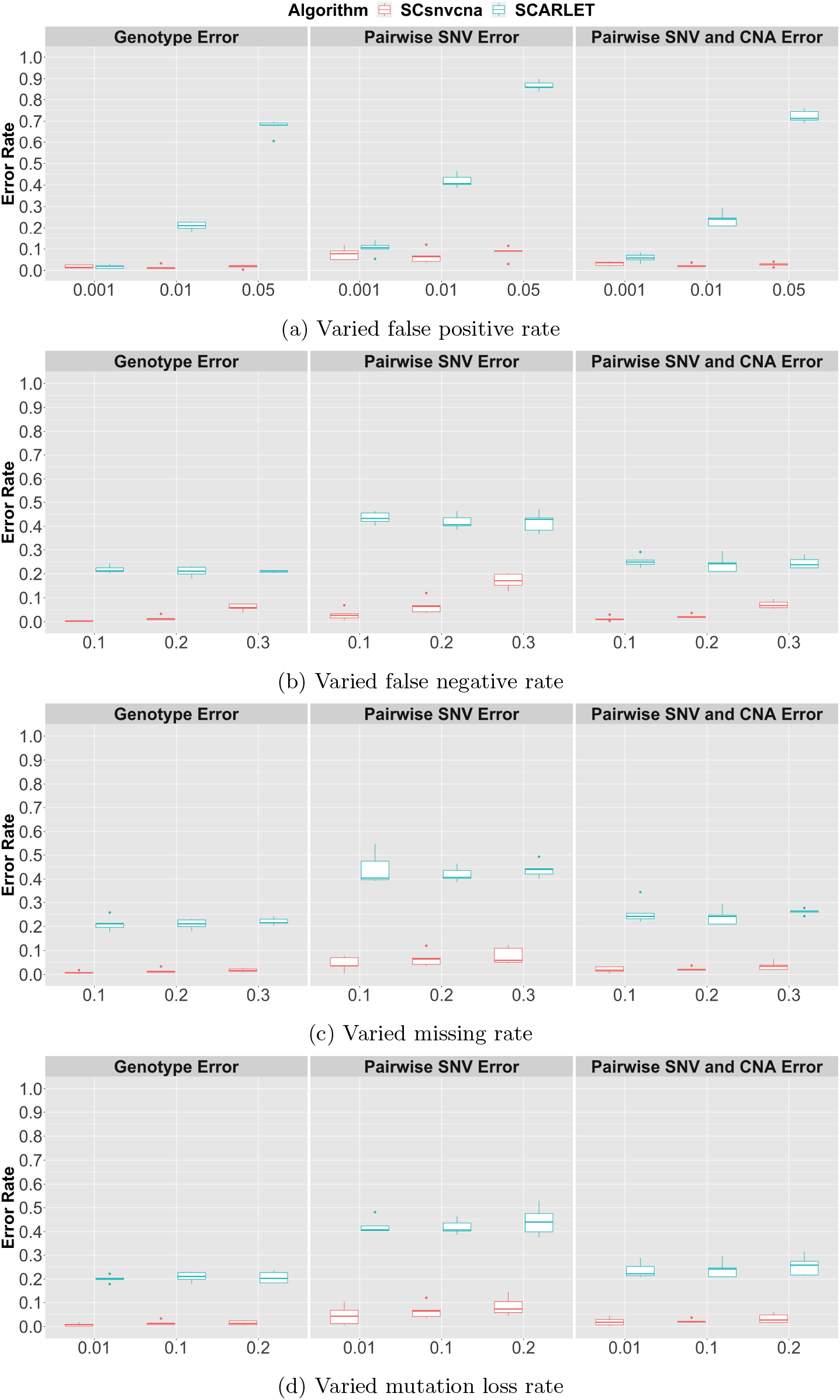
Evaluation of varied parameters

**Fig. 4:**
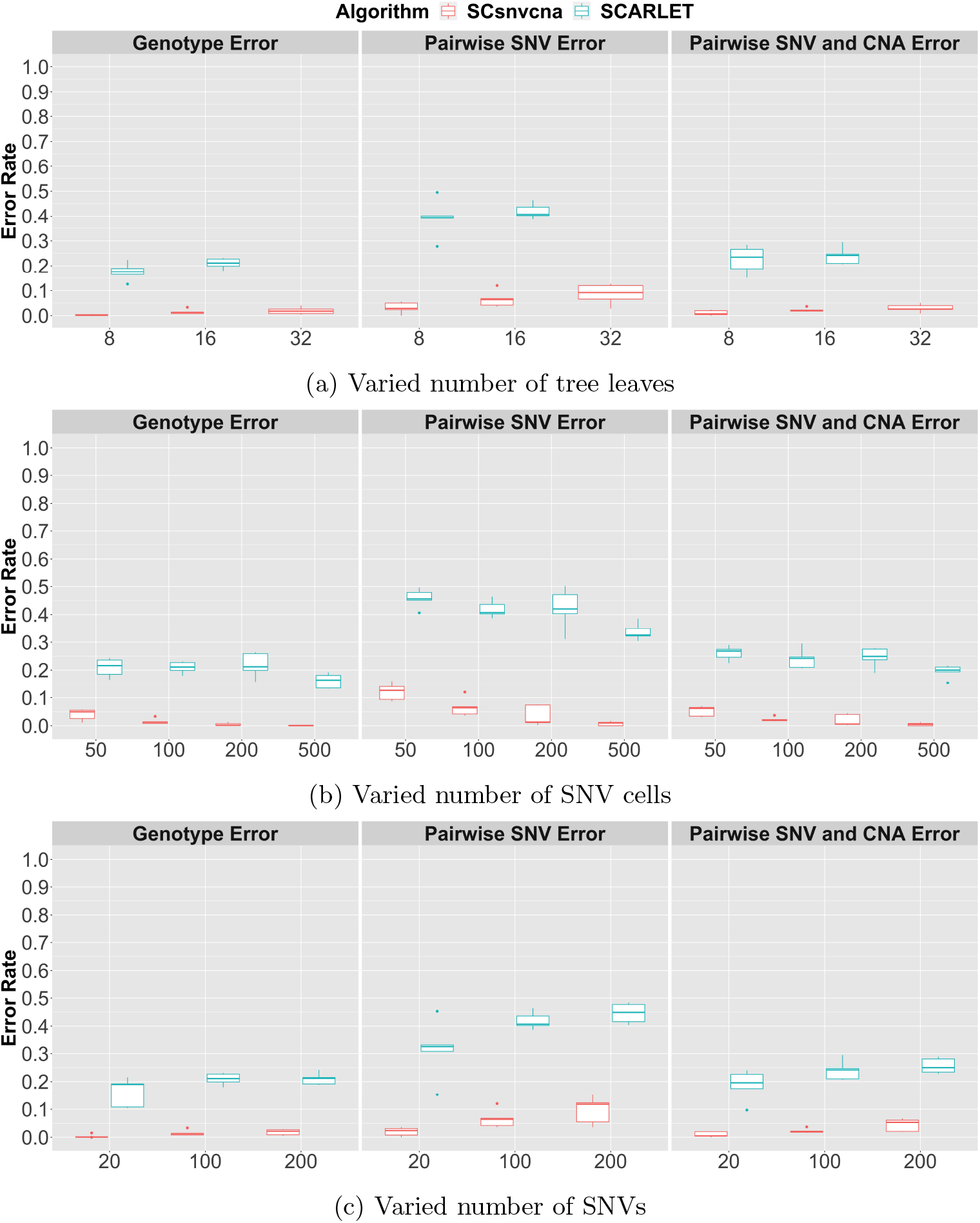
Evaluation of scalability

For all nine variables, SCsnvcna’s genotype error rate stayed lower than that of SCARLET’s, the difference of which was about 0.2 for most of the datasets. We noticed that unlike SCsnvcna which was robust to the false positive rate, SCARLET was very sensitive to high false positive rate (**Fig. 3a**). When false positive error rate was as high as 0.05, SCARLET’s genotype error rate increased close to 0.7; its pairwise SNV error rate was above 0.85 and the pairwise SNV and CNA error rate was above 0.7. When we increased false negative error rate, we noticed a slight increase of genotype error rate, an increase of pairwise SNV error rate and a slight increase of the pairwise SNV and CNA error rate for SCsnvcna (**Fig. 3b**). But even when false negative error rate was as high as 0.3, SCsnvcna’s genotype error rate remained much lower than that of SCARLET’s, by at least 0.1. Similarly, SCsnvcna’s pairwise SNV error rate and pairwise SNV and CNA error rate were about 0.2 lower than that of SCARLET’s when false negative error rate was 0.3. Both SCsnvcna and SCARLET showed a slight increase of the errors with the increasing missing rate, but the increase was very small (**Fig. 3c**). Both SCsnvcna and SCARLET were robust to the increasing mutation loss rate despite a slight increase of pairwise SNV error for both methods (**Fig. 3d**). Similar to what was observed in the varying false positive, false negative and missing rate, SCsnvcna’s genotype error rate, pairwise SNV error rate and pairwise SNV and CNA error rate remained much lower than those of SCARLET for varying mutation loss rates. Specifically, for all mutation loss rates, SCsnvcna’s genotype error rate and pairwise SNV and CNA error rate were around 0.2 lower than those of SCARLET, respectively. Moreover, SCsnvcna’s pairwise SNV error rate was around 0.4 lower than that of SCARLET.

In conclusion, when false positive error rate is as high as 0.05, SCARLET’s genotype result is not reliable since its error rate is as high as 0.7. SCsnvcna is more sensitive to false negative error rate, but even when the false negative error rate is as high as 0.3, its genotyping error rate is still within 0.1 and is at least 0.1 smaller than that of SCARLET’s. Thus SCsnvnca is much more advantageous than SCARLET in the case of high false positive and high false negative error rates.

In the event of increasing subclones which were represented by leaf nodes, we observed that both SCsnvcna and SCARLET’s genotype error rate increased (**Fig. 4a**). However, SCsnvcna’s genotype error rate remained about 0.2 lower than that of SCARLET’s. SCARLET failed to give a solution when the leaf number was as high as 32 whereas SCsnvcna was scalable to such large trees, showing that SCsnvcna is more scalable than SCARLET. Moreover, SCsnvcna’s genotype error rate remained low in the case of large trees, as low as *<* 0.05. We then examined SCsnvcna’s performance under the increasing number of SNV cells and increasing number of SNVs (**Fig. 4b**). We observed that both SCsnvcna and SCARLET’s genotype error rate, pairwise SNV error rate and pairwise SNV and CNA error rate decreased with the increasing number of SNV cells. An opposite trend was observed in the increasing number of SNVs (**Fig. 4c**). We reasoned that when the number of SNV is fixed, more SNV cells provides more constraints on the placement of SNVs, thus help to overcome the errors in the data. On the other hand, when SNV cell number is fixed, more SNVs pose more challenges due to that the search space is larger. Nevertheless, for all the number of SNV cells and number of SNVs, SCsnvcna’s genotype error rate, pairwise SNV error rate and pairwise SNV and CNA error rate stayed much lower than those of SCARLET’s, showing its advantage over SCARLET over a wide range of SNV cell numbers and SNV numbers.

In conclusion, unlike SCARLET, SCsnvcna has been shown to be scalable to large number of leaf nodes and large number of SNV cells and SNVs. Specifically, SCARLET was not scalable to a CNA tree with more than 16 leaves whereas SCsnvcna continued to render low genotype error results even when the number of leaves was as high as 32.

We also measured the performance of SCsnvcna when we varied the variance of CPs between SNV cells and CNA cells (*σ*′), the results of which were shown in **Fig.S2a**. We noticed the genotype error was the lowest when *σ*′ was closest to our initialization (0.05). When *σ*′ was at a different value (0.01 and 0.1), SCsvncna’s genotype error rate remained within 0.05, to be compared with SCARLET’s which were around 0.2 for all values of *σ*′. Lastly, we examined SCsnvcna’s performance when the variance of subclonal sizes varied. This was done by varying the Beta splitting variable described in **Supplementary Section 1**. When Beta splitting variable was close to 0.5, the subclones’ sizes were close to each other, making it hard to identify which edge to place a SNV by the CP. Thus there could be multiple equally good solutions as summarized in [17]. As was expected, the larger the Beta splitting variable, the higher the genotype error rate, pairwise SNV error rate and pairwise SNV and CNA error rate for both SCsnvcna and SCARLET (**Fig.S2b**). However, the increase was very slight and SCsnvcna’s error rates remained much lower than those of SCARLET’s.

In addition to the genotype error rate, pairwise SNV error rate and pairwise SNV and CNA error rate, we also examined the accuracy of SNV loss detection and compared the sensitivity and specificity with that of SCARLET’s for all nine variables (**Fig.S3**). Although SCsnvcna’s sensitivity of SNV loss detection was not as high as SCARLET’s, its specificity was higher than that of SCARLET’s. In fact, its specificity stayed between 0.9 and 1.0 for almost all nine variables. Thus we concluded that SCsnvcna is more conservative in predicting SNV losses than SCARLET.

In all of the above experiments, we fixed SNV cells’ placement for SCsnvcna to conduct and fair comparison with SCARLET. To show that SCsnvcna can achieve a high accuracy even without fixing SNV cells’ placement, we further examined its performance in terms of varying percentage of CNAs that are detectable by SNV cells.

The error rate decreased with the increase of the percentage of CNAs detectable by SNV cells (**Fig. 5**). Genotype error rate was as low as 0.025 when the percentage of CNAs detectable by SNV cells was high (20%). When the percentage of CNAs detectable by SNV cells decreased to 5%, the genotype error rate slightly increased, to be 0.05. Similar patterns were found in pairwise SNV errors and pairwise SNV and CNA errors, benefiting from the fact that a higher percentage of CNAs detectable by SNV cells provided more information on SNV cell placements, and hence lowered the error rate. The evaluation of mutation loss with different levels of percentage of CNAs detectable by SNV cells are shown in **Fig.S4**. We found that SCsnvcna was conservative in predicting SNV losses, leading to a high specificity but low sensitivity in predicting SNV losses. We leave the improvement of sensitivity of SNV loss detection as a future work and discuss it in on Discussion section.

**Fig. 5:**
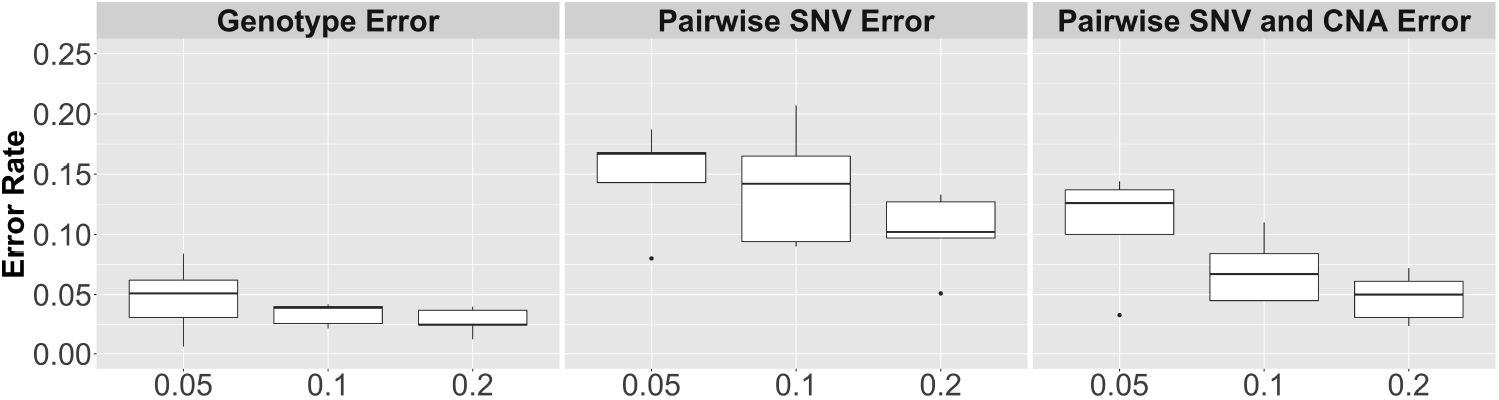
Varied percentage of CNAs detectable by SNV cells

### Real Data Experiment: CRC2

#### Applying SCsnvcna to CRC2

The scDNA-seq data on metastatic colorectal cancer, CRC2 [15], composed of a primary tumor sample and two metastatic samples, serves perfectly the purpose of this study since it contains both CNA cells and SNV cells from the same patient sampled at the same time. In addition, there have been controversial conclusions regarding whether there have been “bridge mutations” in between the two metastasis, i.e., whether the primary tumor continued to gain more mutations after the first metastasis occurred and reseeded that led to the second metastasis. We therefore are interested in applying SCsnvcna on CRC2 to investigate the data by considering both CNAs and SNVs on the same phylogenetic tree. Since the sample has sequencing data from both the primary colon tumor and two metastasis tumors, we used the aneuploid SNV and CNA cells from all three sites.

In more detail, thirty-three CNA cells from the primary colorectal/colon tumor site were used in our CNA analysis. Sixty-seven SNV cells and thirty SNV sites were included as our input to SCsnvcna. To generate a CNA tree as the input to SCsnvcna, we first aligned the reads from CNA cells to the reference genome (GRCh37) by bwa 0.7.17 [16] and detected CNAs by Ginkgo [9] using 0.5Mbp as the bin width. We then masked the bad bins to remove bins with extremely high or low read counts. We found that some copy number profiles inferred by Ginkgo had wrong ploidy, and the baseline upon which the absolute copy number of the whole genome was inferred was either elevated or decreased. This is not an uncommon challenge and has been reported in the previous studies [18,19]. To obtain the correct absolute copy number profiles, we first manually found the baseline of each cell by comparing the whole genome copy number profiles with that of Figure 5D in [15]. In particular, for each copy number profile, we found the copy number corresponding to 2 in [15], which will be used as the baseline for this copy number profile. For those cells whose copy number baselines are greater than 2, we corrected their copy number profiles by finding all those regions whose inferred absolute copy number was larger than the elevated baseline, and deemed these regions as copy number gain regions. For those regions whose inferred absolute copy number was smaller than the elevated baseline, we deemed them as copy number loss region. Those regions whose inferred absolute copy number was the same as the elevated baseline were deemed as copy number neutral regions. In this way, we manually corrected the ploidy for the cells with elevated baselines. The original copy number profiles inferred by Ginkgo for these cells are shown in **Fig.S5**. Notice that such a correction led to a loss of resolution on the absolute copy number because we will no longer have the numbers but just three categories, copy number gain/loss/neutral.

The baseline correction for the cells whose baseline was lower than two was different from those with elevated baselines due to that the bona fide copy number loss regions and copy number neutral regions could have been merged and thus no longer be separated. We therefore did not use these cells to infer the CNA tree. Such cells include MA55, MA58, MA60, MA64, MA67, MA68, MA69, MA70 and MA71, and their copy number profiles are shown in **Fig.S6**.

We then inferred a maximum parsimony tree using PAUP [28] from the corrected copy number profiles, where the genome at the root of the tree was assumed to be a diploid cell without any CNAs.

Notice that we used the CNA cells from all three tumors (primary, metastasis 1 called M1 and metastasis 2 called M2) of the same patient sample for inferring the CNA tree without fixing their genealogical order on the tree. This means primary cells may be in a parallel branch of the metastasis cells or even occur later than the metastasis cells. Since the diagnosis of the primary tumor and the two metastatic tumors in CRC2 occurred at the same time [15], the primary tumor cells may have evolved after the metastasis happened, as also recognized in [15]. Therefore instead of assuming that all primary cells stay closer to the root than M1 and M2, we used the maximum parsimony method to infer the CNA tree based on all CNA cells in order to best explain the lineage of the primary cells and the two sets metastatic cells in CRC2.

In preparing the *D* matrix as the other input to SCsnvcna, we obtained it from [15] and eliminated those SNVs whose CP is *<* 5%. These SNVs were not used due to the limited resolution provided by CNA cells as there were only 33 CNA cells being used in the CNA tree.

We then identified the potential mutation loss list by pairing up overlapping copy number losses and SNVs. Any SNVs overlapping with these copy number losses and were placed on the path between the root and the edges that the copy number losses were placed can potentially be lost, and we left the decision whether the SNV is lost to the SCsnvcna’s algorithm.

The last step in preparation for running SCsnvcna was to identify whether it was possible to nail down some SNV cells’ placement on the CNA tree by the CNAs observable from the read depth in these SNV cells. To perform this task, we inferred the copy number profile for each SNV cell by Ginkgo [9] using the same parameter setting as that of the CNA cells. While it was reasonable to use a larger window size for SNV cells due to its non-uniform read depth, we chose to use the same window size (0.5Mbp) not only because this window size was in the middle of those fitting for both CNA cells and SNV cells, but also because we aimed at achieving the same resolution of CNAs from both CNA cells and SNV cells. Not surprisingly, like CNA cells, we again observed that a large percentage of the SNV cells’ baselines of the copy number profile were either elevated or decreased, showing that Ginkgo failed to correctly infer the copy number profiles’ baselines for SNV cells. We thus manually corrected the baselines of SNV cells’ copy number profiles in a similar manner as we did for the CNA cells’, and simplified the absolute copy number profiles to be just three categories, copy number gain, loss and neutral. We then adopted the second method as described in the section “Using the CNAs on SNV cells to limit the placement of SNV cells” to nail down some SNV cells’ placement on the CNA tree. In more detail, we compared pairwise copy number profiles from SNV cells and CNA cells and computed the percentage of CNA segments that have the same copy number event for each pair. A SNV cell’s placement will be constrained to a leaf node if it shares ≥ 80% copy number gain/loss events with those of the CNA cells this leaf node carries. If there were more than one leaf node that the SNV cell should be placed, we left the decision to SCsnvcna according to whichever leaf node would lead to the highest joint probability.

Since this dataset’s false positive and false negative rates have been estimated in [15], we used the estimated values, 1.74% and 12.56%, respectively, to initialize *α*_*a*_, *α*_*b*_, *β*_*a*_ and *β*_*b*_. We then ran SCsnvcna by the default setting on the CRC2 data.

#### CRC2 Results

A simplified phylogenetic tree as the output of SCsnvcna on CRC2 was shown in **Fig. 6**. A detailed phylogenetic tree with the labels of the single cells was shown in **Fig.S7**.

**Fig. 6:**
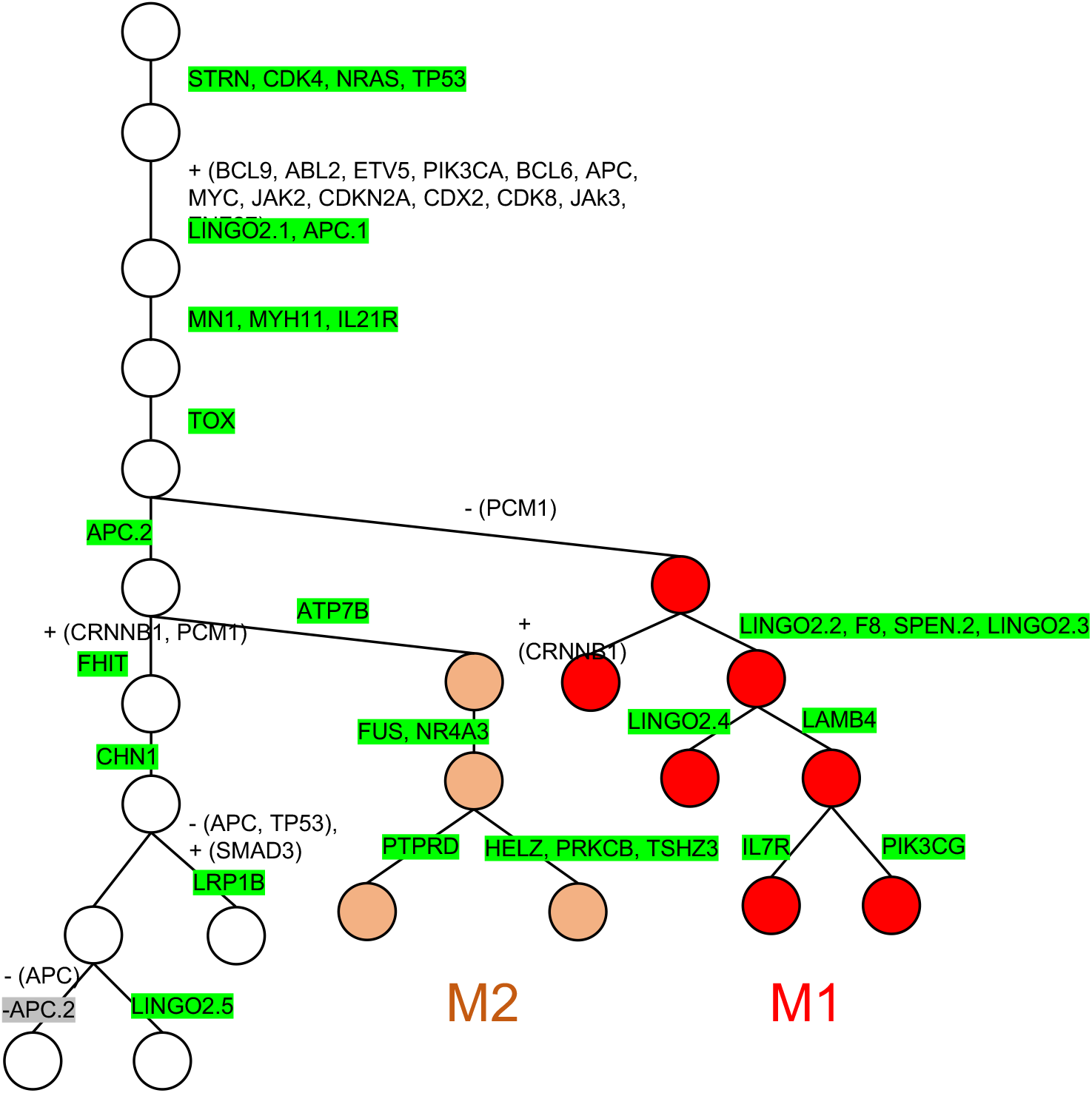
A simplified phylogenetic tree for CRC2 inferred by SCsnvcna. Circles represent the tumor clones. Metastatic cells belonging to M1 and M2 are highlighted in red and orange, respectively, whereas all other nodes represent cells in the primary tumor. The genes of CNAs and SNVs are annotated on the edges of the tree. SNVs are highlighted with the green background. CNAs are in the white background. Copy number gains and losses are grouped within parenthesis and denoted by a “+” and “-” sign, respectively. SNV losses due to copy number loss are highlighted by the gray background denoted with a “-” sign.

Like Leung et al., SCsvncna placed the major SNVs and CNAs on the trunk. Specifically, SCsnvcna placed the TP53 mutation before the activation of oncogenes BCL9, ABL2, ETV5, PIK3CA, BCL6, MYC, JAK2, CDKN2A, CDX2 and CDK8 through copy number gains. SCsnvcna then further placed mutations on MN1, MYH11 and IL21R on the trunk, followed by a mutation on TOX, another tumor suppressor gene. The tumor then diverged and the first metastasis (M1) occurred after a copy number loss on PCM1. The other branch gained a mutation on the tumor suppressor gene APC, and further branched into the second metastasis (M2) after gaining a mutation on ATP7B.

On the cluster of M1, SCsnvcna and Leung et al. [15] were consistent in placing the SNVs of LINGO2.3, F8, SPEN, LINGO2.2, LAMB4, IL7R, PIK3CG and LINGO2.1. The only difference on M1 was that Leung et al. [15] placed the SNV of PTPRD also on M1 whereas SCsnvcna placed it on M2. We noticed out of thirteen cells having the mutation signal on PTPRD, Leung et al. assigned only three cells as true positives. Similarly, SCsnvcna also assigned three out of thirteen as true positives, although these three cells (MA34, MA87, MA95) were different from the three cells that Leung et al. deemed as true positives (MA86, MA35, MA33). Thus whether PTPRD occurred in M1 or M2 is uncertain.

On the cluster of M2, SCsnvcna was consistent with Leung et al. [15] in placing the SNVs of FUS, NR4A3, HELZ, PRKCB and TSHZ3. Similar to Leung et al. [15], SCsnvcna placed ATP7B on top of all M2 cells, confirming that ATP7B was the mutation that initiated the second metastasis in liver. This is consistent with the study of ATP7B [23] that showed ATP7B is highly expressed in liver and intestine and the mutations on ATP7B may lead to symptoms of Wilson disease in both liver and intestine [23].

Both SCsnvcna and Leung et al. inferred that after M1 occurred, the primary tumor cells continued to gain more mutations on genes CHN1, FHIT, LINGO2 and LRP1B (listed as UAP1B in Leung et al. [15]). Like Leung et. al [15], SCsnvcna also inferred a bridge mutation on APC that occurred on primary cells after M1 but before M2. This mutation was then inherited by all the cells in M2. This conclusion is different from what SCARLET [26] indicated, which placed the mutation of APC above M1, and inferred that it was lost due to a copy number loss on the branch leading to M1. While there were copy number losses occurring to APC, they mostly occurred on the branch of primary cells near the leaves. SCsnvcna did not report a copy number loss on APC on the cluster of M1. Neither did it report a loss of mutation of APC on M2. SCsnvcna, however, inferred that a few primary SNV cells lost the mutation of APC due to the copy number loss on APC. Based on our simulation study which showed a high specificity in predicting mutation loss due to copy number loss, such a prediction of mutation loss on APC of the primary cells is likely to be true.

We then investigated SCsnvcna’s placement of SNV cells by comparing their clustering with the anatomical origin from the patient. We found that SCsnvcna clustered the SNV cells into three main clusters: primary, M1 and M2 and thus was consistent with the resection sites of the cells. Specifically, thirteen out of sixteen cells that were clustered by SCsnvcna have been categorized as M1 in [15]. SCsnvcna clustered all thirteen SNV cells that were classified as M2 in [15].

## Discussion

We developed SCsnvcna, an open source computational tool that places SNVs and SNV cells on a CNA tree for scDNAseq data. The algorithm is based on the fact that the SNV cells and CNA cells from the same subclone shall have close cellular prevalence. SCsnvcna takes into account the scDNAseq-specific error profile such as false positive and false negative rates, the missing errors, as well as the potential loss of SNVs due to copy number losses. Moreover, SCsnvcna can utilize the limited CNA profiles indicated by the SNV cells to constrain the SNV cell placement on a CNA tree. Finally, SCsnvcna was designed to model and thus tolerate the sampling bias between CNA and SNV cells.

Nine simulated datasets have been used to examine the performance of SCsnvcna in comparison with SCARLET [26]. SCsnvcna’s genotype error stayed about 0.2 lower than that of SCARLET’s for almost all datasets. Moreover, SCsnvcna was more scalable than SCARLET. While SCARLET failed to render a solution when the number of leaf number was as high as 32, SCsnvcna continued to render a solution which had low genotype error rate. We applied SCsnvcna to CRC2 dataset and found that SCsnvcna’s placement of SNVs was consistent with that of Leung et al [15]. SCsnvcna also correctly placed most of the cells which clustered into primary and two metastasis groups according to their anatomical sites. Interestingly, SCsnvcna inferred a “bridge” SNV at APC between the two metastases which was consistent with the conclusion from the original study on CRC2 in [15], thus confirmed that there was a bridging and reseeding process after the first metastasis was formed.

One future work of improving SCsnvcna is to enhance its sensitivity of mutation loss detection. Specifically, for those mutation losses caused by copy number losses, comparing the read depth on the locus where the mutation losses occur with its flanking regions will help to identify potential mutation losses.

Another interesting route as a future work is to try to place CNAs on a SNV tree, a tree whose structure is decided by SNVs. We did not seek this route due to that the CNA tree provides us the information of copy number losses which can guide the placement of SNVs in case of mutation losses. A third route is to jointly infer the phylogenetic tree, the placement of SNVs and CNAs, as well as the placement of SNV cells and CNA cells simultaneously. Although this route may provide more accurate solutions since the phylogenetic tree is inferred based on both SNVs and CNAs, it involves a much bigger solution dimension and thus is computationally challenging. We leave this route to a future work as an improvement to SCsnvcna.

In conclusion, SCsnvcna is a pioneer computational phylogenetics tool that solves the challenge of integrating previously disparate single-cell datasets by mapping SNVs onto a CNA tree. SCsnvcna can validate and reveal interesting order and interplay between SNVs and CNAs and thus boost the understanding of how tumor gains new mutations and grows. With the decease of the sequencing cost, SCsnvcna will be widely applied to multiple scDNAseq data sets and help to understand tumor growth and thus resolve ITH.

## Acknowledgement

X.M. was supported by the startup funding from Florida State University. L.Z. was supported by the Planning Grant and Small Grant from the The Council on Research & Creativity of Florida State University.

## Data and Code Availability

SCsnvcna software as well as the simulator described in this study are publicly available at https://github.com/compbio-mallory/SCsnvcna. CRC2 raw data was downloaded from NCBI Sequence Read Archive under accession number SRP074289.

